# Adult-specific trimethylation of histone H3 lysine 4 is prone to dynamic changes with aging in *C. elegans* somatic cells

**DOI:** 10.1101/236257

**Authors:** Mintie Pu, Minghui Wang, Wenke Wang, Satheeja Santhi Velayudhan, Siu Sylvia Lee

## Abstract

Tri-methylation on histone H3 lysine 4 (H3K4me3) is associated with active gene expression but its regulatory role in transcriptional activation is unclear. Here we used *Caenorhabditis elegans* to investigate the connection between H3K4me3 and gene expression regulation during aging. We uncovered around 30% of H3K4me3 enriched regions to show significant and reproducible changes with age. We further showed that these age-dynamic H3K4me3 regions largely mark gene-bodies and are acquired during adult stages. We found that these adult-specific age-dynamic H3K4me3 regions are correlated with gene expression changes with age. In contrast, H3K4me3 marking established during developmental stages remained largely stable with age, even when the H3K4me3 associated genes exhibited RNA expression changes during aging. Moreover, we found that global reduction of H3K4me3 levels results in significantly decreased RNA expression of genes that acquire H3K4me3 marking in their gene-bodies during adult stage, suggesting that altered H3K4me3 levels with age could result in age-dependent gene expression changes. Interestingly, the genes with dynamic changes in H3K4me3 and RNA levels with age are enriched for those involved in fatty acid metabolism, oxidation-reduction, and stress response. Therefore, our findings revealed divergent roles of H3K4me3 in gene expression regulation during aging, with important implications on physiological relevance.

## Introduction

Aging in diverse organisms are accompanied by alteration in gene expression profiles that tightly correlate with age-dependent physiological changes [1-13]. A recent study in *C. elegans* showed that the precise regulation of gene expression plays an important and causal role in the aging process, with inhibition of aging-dependent gene expression drift extends lifespan [14].

Epigenetic mechanisms are key to gene expression regulation [15]. Previous studies demonstrated that dynamic changes in DNA méthylation, histone modifications, and small RNAs levels occur during aging in different species [16-22]. However it remains unclear how the age-dependent changes in various epigenetic profiles relate to physiological changes.

Tri-methylation on histone H3 lysine 4 (H3K4me3) is a widely recognized active promoter mark [23]. It was reported that over 80% of the genes marked with H3K4me3 at the promoter region are actively transcribed [24]. H3K4me3 has been implicated in regulating preinitiation complex formation and gene activation [25], and in regulating pre-mRNA splicing [26], DNA recombination [27], DNA repair [28] and enhancer usage [29]. However, studies in yeast and mammalian cells showed that defects in major H3K4me3 histone modification enzymes do not result in changes in gene expression [30-32], indicating that loss of H3K4me3 has minor effects on global transcription. These data suggest that H3K4me3 could be deposited after gene activation and represents a record of gene expression history. Most recently, studies in mammals showed that broad H3K4me3 regions function in maintaining cell identity [33] and are essential for the expression of tumor suppressor genes [34].

In *C. elegans,* H3K4me3 modulates lifespan in a germline-dependent manner. Inactivation of the major H3K4me3 methyltransferase complex results in extended lifespan, partly through altered fat metabolism [35,36] [37]. In contrast, reduced expression of H3K4me3 demethylases shortens lifespan [35,38]. H3K4me3 also functions in germline development as inactivation of the H3K4me3 methyltransferase and demethylase leads to fertility defects [39,40]. How regulation of H3K4me3 deposition could mediate the various physiological changes remain largely unknown.

In this study, we investigated the connection between H3K4me3 and gene expression regulation in the context of aging. We profiled the dynamic changes of H3K4me3 in *C. elegans* somatic cells at aging time points. We found that around 30% of the H3K4me3 enriched regions exhibit significant changes in H3K4me3 levels with age. The data showed that these age-dynamic H3K4me3 regions preferentially mark gene-bodies and are acquired during adult stages. When compared with parallel RNA-seq analyses, we found that the age-dependent changes of H3K4me3 are generally positively correlated with gene expression dynamics during aging. In particular, the H3K4me3 marks specifically deposited in adulthood were more prone to change with age and coupled with gene expression changes. In contrast, H3K4me3 marking established during developmental stages remained largely stable with age, even when the H3K4me3 associated genes exhibited RNA expression changes during aging. Moreover, our analysis showed that global reduction of H3K4me3 through RNAi of *ash-2,* which encodes a major H3K4me3 methyltransferase complex component, results in the specific decreased expression of genes that acquire H3K4me3 marking during the adult stage. Taken together, the data suggested that the age-dependent altered levels of H3K4me3 could regulate gene expression during aging. The findings reported here revealed two divergent roles of H3K4me3 in gene regulation with aging: The H3K4me3 markings established during development that remain stable with age and likely do not regulate gene expression changes, and the H3K4me3 markings deposited during adult stage that tightly correlate with gene expression changes with age. Gene ontology (GO) term analysis showed that the genes with H3K4me3 and RNA expression changes during aging are enriched those participating in fatty acid metabolism, oxidation-reduction, and stress response, potentially providing a mechanistic understanding of the link between H3K4me3 dynamics and physiological changes during the aging process.

## Results

### Identification of age-dependent H3K4me3 changes in *C. elegans* somatic cells

We performed chromatin immunoprecipitation coupled with deep sequencing (ChlP-seq) to profile the genome-wide patterns of H3K4me3 in *C. elegans* somatic cells at the young and old age (SI Table). We used whole worm extracts from the temperature-sensitive *glp-l(e2141ts)* mutant that produces very few germ cells at the non-permissive temperature[41] to avoid the interference from the germline. We examined three independent biological replicates at two time points: day 2 (D2) adults, a time when wild-type worms are highly reproductive, as the young stage point, and D12 adults, a time when ~10% of the population has started to die, as the old stage point.

To compare the H3K4me3 profiles between young and old stages, we first performed a genome-wide correlation analysis. Normalized H3K4me3 levels were computed using H3 ChlP-seq data as a control. Pearson correlation analysis was performed using normalized H3K4me3 in 2 kb windows tiling across the genome (Fig 1A). We found that the genome-wide pattern of H3K4me3 did not drastically change with age, as the overall correlation coefficients were all higher than 0.74. Nevertheless, the clustering analysis revealed that replicates from the same time point were clustered closer together, suggesting noticeable time point differences. We further used MACS2 (2.1.0) to identify H3K4me3 enriched regions at each time point (S2 Table). Principal component analysis (PCA) was used to compare the H3K4me3 peaks identified from these 2 time points based on their covariance (Fig 1B and S1A Fig). The PCA plot showed that the H3K4me3 enriched regions of D2 adults clearly separated from those of D12 adults along PC2, whereas the replicates for each time point only marginally separated from each other along PCI (Fig 1B and S1A Fig), supporting our earlier conclusion that the H3K4me3 marking at young and old age shows detectable differences. Scatter plot analysis of the normalized H3K4me3 levels for each peak at the D2 and D12 time points also clearly revealed peaks that showed either increased or decreased with age (S1B Fig).

**Figure 1.**
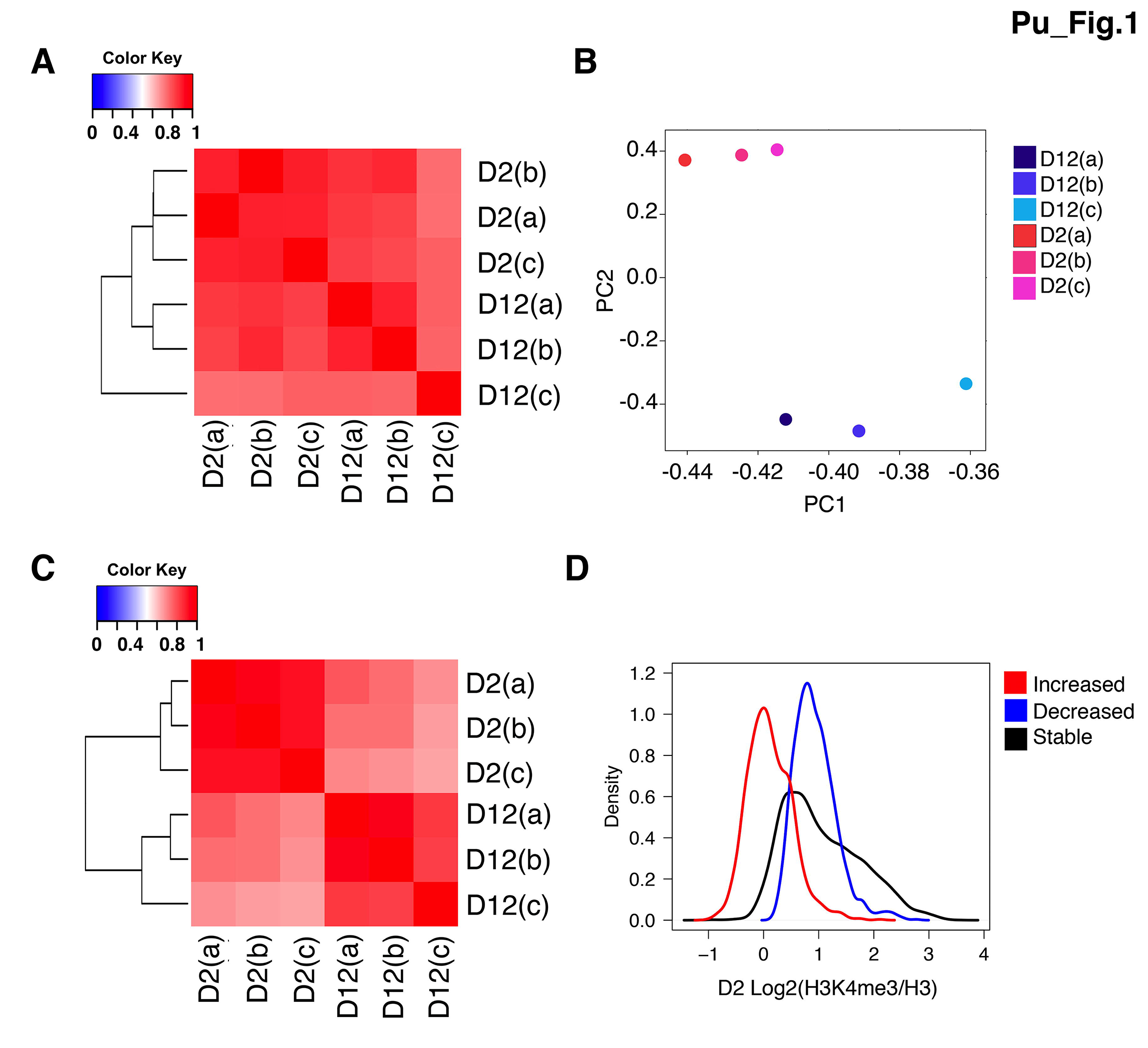
H3K4me3 profiles dynamically change with age in *C. elegans* somatic cells. (A) Genome-wide correlation analysis of H3K4me3 profiles in D2 (young) and D12 (old) *C. elegans.* Pair-wise Pearson correlations of genome-wide H3K4me3 levels were calculated using 2kb windows. (B) PCA plot showing normalized H3K4me3 data from three biological replicates. The H3K4me3 peak regions in D2 and D12 were determined using the MACS2 narrow peak calling method. (C) Correlation analysis of the peak regions that showed significant changes with age as identified by Diffbind. Normalized H3K4me3 levels were used for Pairwise Pearson correlation analysis. (D) Density plot showing normalized H3K4me3 levels at D2 for peaks that showed increased (red), decreased (blue) or stable (black) modification levels with age.

To further identify the specific regions with significant age-dependent H3K4me3 change, we used the Diffbind (version 3.2.1)[42] statistical tool to perform differential analysis, and identified 2,568 differential regions (S3 Table). Pearson correlation analysis of the normalized H3K4me3 signals within these differential regions confirmed that there were obvious differences between young and old stages (Fig 1C). Density plots of normalized H3K4me3 levels for the peaks that increased (1,168), decreased (1,400), or remained stable (5,152) with age showed that the H3K4me3 peaks that increased with age tend to be marked by low levels of H3K4me3 at D2, and the H3K4me3 peaks that decreased with age tend to be marked by medium levels of H3K4me3 at D2 (Fig 1D). The peaks marked with high levels of H3K4me3 at D2 tend to remain stable with age.

We next assigned the age dynamic H3K4me3 peaks to the closest annotated genes (S4 Table). The majority of the dynamic H3K4me3 peaks overlapped or spread into gene body regions (2,190 peaks), whereas a small proportion of peaks (378) located in intergenic regions and were assigned its closest downstream gene. 837 peaks overlapped with more than one gene and were assigned to more than one gene (S4 Table). As expected, the majority of the dynamic H3K4me3 peaks were assigned to protein coding genes (S1C Fig).

### H3K4me3 markings that span gene body are significantly enriched for age-dependent H3K4me3 changes

Since the majority of the dynamic H3K4me3 peaks were assigned to protein coding genes (S1C Fig), we focused our downstream analyses using only the H3K4me3 peaks that were assigned to protein coding genes. To identify any possible patterns associated with the age-dependent dynamic H3K4me3 peaks in an unbiased manner, we performed K-mean clustering of the 9,213 protein-coding genes associated with H3K4me3 peaks (S5 Table). Normalized H3K4me3 levels spanning each gene together with 2kb upstream and downstream regions at the D2 and D12 time points were used for the clustering analysis. Empirical trials identified 25 clusters that represented the diversity of the H3K4me3 pattern for these genes (Fig 2A). The heatmap revealed that, in addition to the typical H3K4me3 pattern that concentrated around annotated transcriptional start sites (TSS) (clusters a-j) (WBcel235), there were also clusters with H3K4me3 signals spreading evenly across gene body regions (clusters 1-q) (Fig 2A).

**Figure 2.**
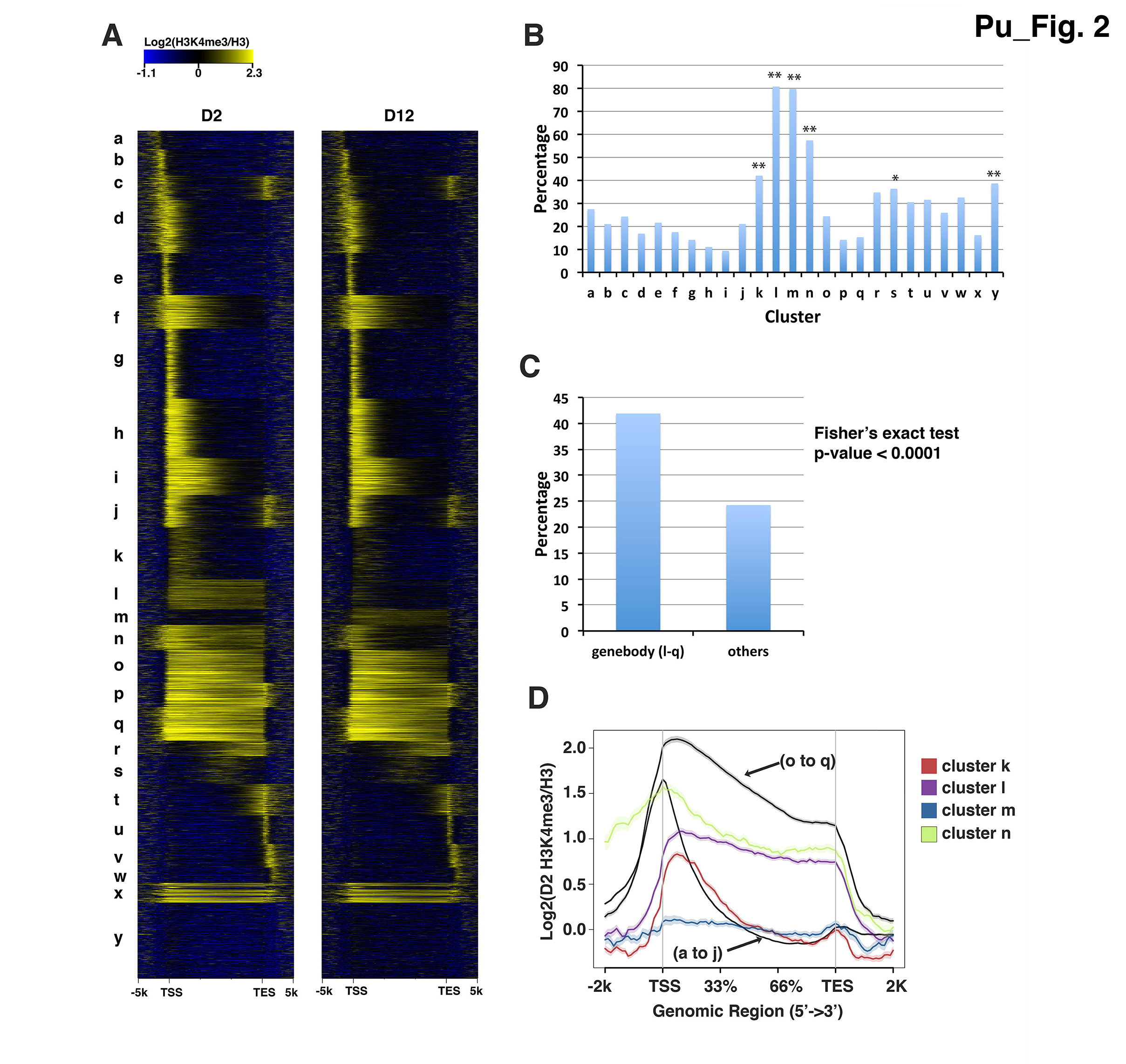
H3K4me3 marking on gene body are enriched for age-dependent H3K4me3 changes. (A) Protein-coding genes associated with H3K4me3 peaks (identified by MACS2) were grouped into 25 clusters (a-y) using k-means clustering based on their normalized H3K4me3 levels at D2 and D12 time points. The heat map shows log2 ratio of H3K4me3 levels normalized to H3 levels. (B) The bar chart shows the percentage of genes associated with age-dependent dynamic H3K4me3 peaks identified by Diffbind for each of the 25 clusters shown in (A). Clusters k, l, m, n, s and y were significantly enriched for genes associated with H3K4me3 peaks that dynamically changed with age. (*), Fisher’s exact test, p-value<0.05. (**) Fisher’s exact test, p-value<0.01. (C) H3K4me3 peaks spanning gene body are enriched for dynamic changes with age. The percentage of genes associated with age-dependent H3K4me3 dynamic change for clusters l to q, where H3K4me3 marking spanned gene body, were compared with that for all the other clusters using Fisher’s exact test. (D) Average plots show the normalized H3K4me3 levels for the indicated clusters. Clusters k, l, m and n, which were enriched for age-dynamic H3K4me3 changes (shown in B), were marked with relatively lower levels of H3K4me3.

Interestingly, the heatmap showed that the H3K4me3 markings that spanned gene body regions appeared enriched for dynamic changes with age, including clusters 1, m, and n (Fig 2A). To further investigate this observation, we located the genes associated with the age-dynamic H3K4me3 peaks identified by Diffbind previously (2,544 genes) into these 25 clusters (S4 Table) and calculated the percentage of genes in each cluster with age-dynamic H3K4me3. As expected, clusters 1, m, and n, which contained genes marked by H3K4me3 throughout their gene bodies, were significantly enriched for age-dependent H3K4me3 changes (Fig 2B). Clusters k and s, which showed weak partial gene body H3K4me3 marking, were also significantly enriched for age-dependent H3K4me3 changes (Fig 2B). Fisher’s exact test confirmed that, as a group, the genes with gene body H3K4me3 marking (clusters 1 to q) were associated with more dynamic age-dependent change of H3K4me3 compared with genes with other H3K4me3 marking pattern (clusters a-k, r-y) (Fig 2C). Furthermore, average plot analysis showed that the H3K4me levels in clusters 1, m, and n, were relatively lower than those of the other gene body clusters ((o-q) (Fig 2D), suggesting that gene body regions marked by low levels of H3K4me3 are particularly prone to dynamic regulation of H3K4me3 with age.

### Age-dynamic H3K4me3 markings are mainly deposited in adult stage

To understand the possible temporal window that generates the different patterns of H3K4me3, we next compared the genome-wide H3K4me3 profiles in larval and adult stages in *C. elegans.* We performed H3K4me3 ChlP-seq in L3 stage germline-less *glp-1* mutant worms and identified 6,104 H3K4me3 peaks (S6 Table). We found that the majority of the H3K4me3 peaks detected in the L3 stage were maintained in adults. We further compared the H3K4me3 profiles at L3, D2 and D12 stages using the 25 clusters generated by K-mean clustering discussed previously (S2A Fig) and computed the average normalized H3K4me3 levels for each cluster at the L3 and D2 time points (Fig 3A). Strikingly, this comparison revealed that clusters k, 1, m and n, which we found to be enriched for genes associated with age-dynamic H3K4me3 peaks (Fig 2), were marked with low or undetectable levels of H3K4me3 at the L3 stage compared to D2 or D12 (Fig 3A, S2A Fig), suggesting that the H3K4me3 markings for these gene clusters were mainly deposited during the adult stage.

**Figure 3.**
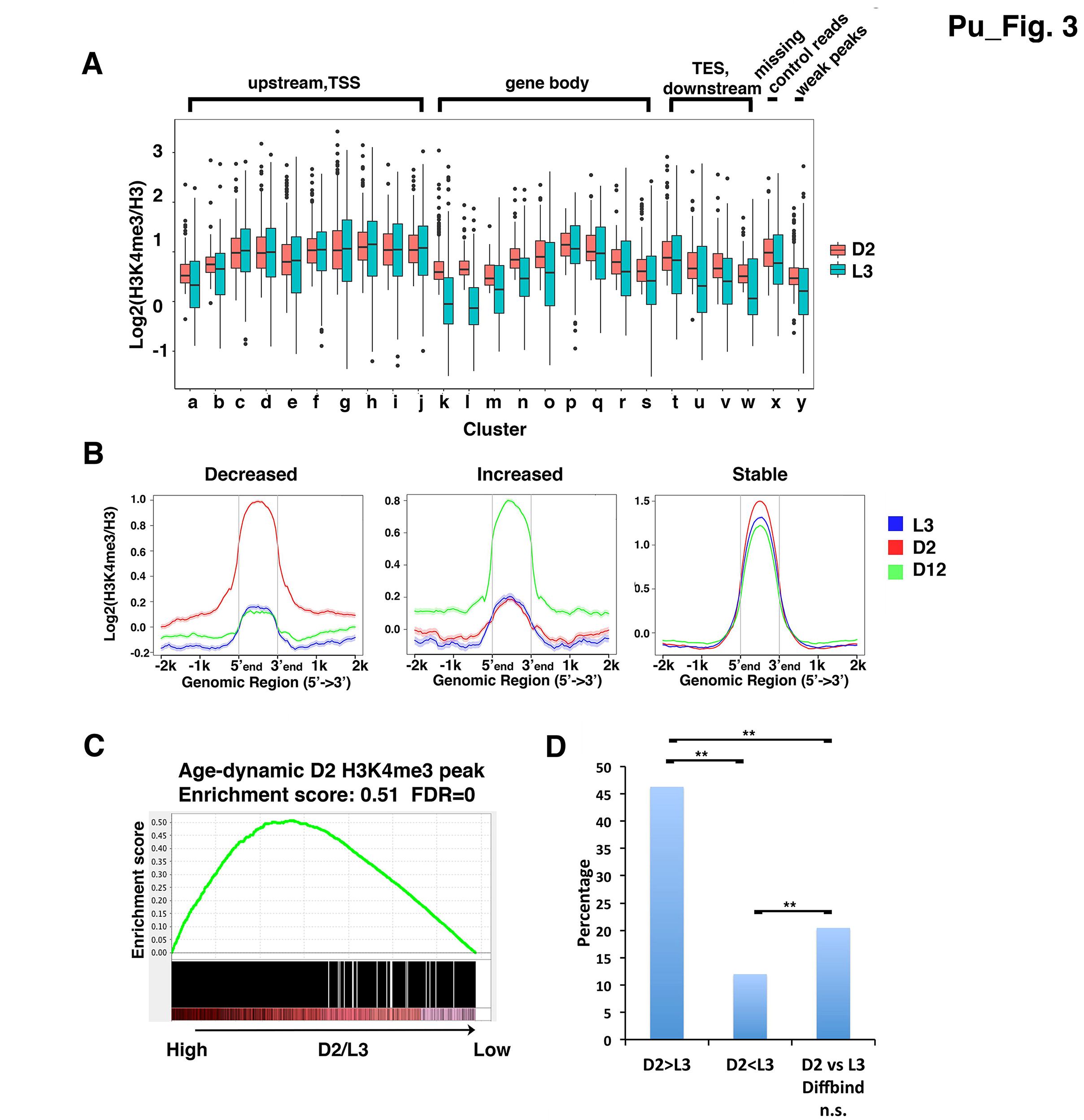
The age-dynamic H3K4me3 markings are mainly deposited in adults. (A) The boxplot shows the normalized H3K4me3 levels in the 25 clusters at D2 or L3 stages. (B) Average plots of the H3K4me3 levels at larval stage 3 (L3) (blue), D2 (red) and D12 (green) for peaks that showed significant dynamic changes with age identified by Diffbind. Normalized H3K4me3 levels within the peak regions and 2kb upstream and downstream are shown for the peaks that showed significant decrease (left), increase (middle), or stable (right) H3K4me3 levels with age. (C) Age-dynamic HK4me3 peaks are enriched for peaks marked with higher levels of H3K4me3 at D2 relative to L3. To perform GSEA (Gene Set Enrichment Analysis), H3K4me3 peaks at D2 & D12 were ranked according to the ratio of H3K4me3 levels at D2 relative to L3. Peaks with a higher ratio were assigned a higher rank and displayed on the left side of the heatmap. Age-dynamic H3K4me3 peaks were extracted and compared with the list of D2/D12 H3K4me3 peaks. The enrichment scores were computed using the GSEAPreranked tool. An enrichment score of 0.51 represents a statistically significant enrichment for peaks with a high D2/L3 H3K4me3 ratio. (D) H3K4me3 deposited during adult stage are enriched for age-dynamic H3K4me3 peaks. Diffbind was used to identify the H3K4me3 peaks with statistically significant differences between D2 and L3 stages. The differential (D2>L3, or D2<L3) or stable (D2 vs L3 ns) peaks were further compared with age-dynamic peaks to identify the overlapping peaks. The bar chart shows the percentage of age-dynamic H3K4me3 peaks for each of the D2/L3 group. (**) Fisher’s exact test, p-value<0.0001.

To further examine the relationship between age-dynamic H3K4me3 regions with the timing of their initial deposition, we computed the average plots of the age-dynamic H3K4me3 peaks identified by Diffbind previously at the L3, D2 and D12 stages (Fig 3B and S3A Fig). The data showed that for both the age-dependent increased or decreased group, the H3K4me3 levels at L3 were significantly lower than that at D2 or D12 stages (Fig 3B). In contrast, for the H3K4me3 peaks that remained stable with age, they were already marked with comparable levels of H3K4me3 at the L3 stage (Fig 3B).

We investigated this observation further using the statistical tool GSEA (Gene Set Enrichment Analysis) [43], which uses rank order to assess bias distribution, to test whether the age-dynamic H3K4me3 peaks were more likely to gain their H3K4me3 marking in the adult stage. The results supported that the age-dynamic H3K4me3 peaks were indeed significantly overrepresented for peaks with higher H3K4me3 marking at D2 compared to L3, indicating that the age-dynamic peaks are enriched for regions gaining H3K4me3 marking during adult stage (Fig 3C).

To analyze this observation in a converse manner, we examined whether the adultstage deposited H3K4me3 regions could be enriched for age-dynamic changes. We used Diffbind to identify the H3K4me3 peaks with significant difference between D2 and L3 (S7 Table). We then compared the overlap between these L3/D2 differential peaks and the age-dynamic D12/D2 H3K4me3 peaks we identified earlier. The results showed that regions marked with significantly higher H3K4me3 levels at D2 stage were greatly enriched for age-dynamic peaks (Fig 3D). The peaks with higher H3K4me3 levels at L3 stage relative to D2 had the least overlap with the age-dynamic peaks (Fig 3D). Taken together, the data suggested that H3K4me3 deposited during developmental stages remain relatively stable during aging, whereas the H3K4me3 marking deposited in adult stage are more prone to change with age.

### Age-dependent H3K4me3 and gene expression changes are highly correlated

To investigate whether age-dynamic H3K4me3 changes correlated with gene expression changes, we performed parallel ribo-minus RNAseq analyses using the germlineless *glp-1* mutant worms at the exact time points used for the H3K4me3 ChlP-seq experiments (S8 Table). The overall correlations between the two biological replicates from each of the time points were both higher than 0.90, indicating the high reproducibility of the experiments. A comparison with our previously published poly-A RNAseq results with a similar experimental design [44] showed that the two RNAseq datasets were highly correlated at both the young and old time points (rho = 0.92 at D2 and D12) (S1 Fig). With the ribo-minus RNAseq data, the statistical tool edgeR identified 3,154 genes that showed significant expression change during aging. Among these, about half of the genes were marked with H3K4me3 (1,902), and among this subgroup, ~50% (919 genes) correlated with age-dependent H3K4me3 changes (Fig 4A). Importantly, the direction of RNA expression change and H3K4me3 change was significantly positively correlated, with a Spearman’s correlation coefficient of 0.65 (Fig 4B, S9 Table).

**Figure 4.**
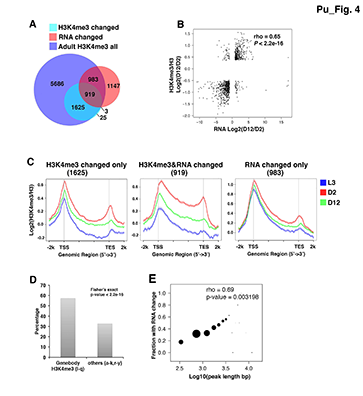
Age-dynamic H3K4me3 peaks accompanied by RNA expression change are more likely to span gene body regions and be deposited during adult-stage. (A) The venn diagram shows the protein-coding genes that were associated with H3K4me3 marking in adult stages (blue), the ones associated with significant H3K4me3 change with age (aqua), and the ones associated with significant RNA expression change with age (red). Gene numbers for each group are shown. (B) Age-dependent H3K4me3 change and gene expression change are positively correlated. Spearman’s correlation coefficient was calculated using log2 ratio of normalized H3K4me3 or RNA levels at D12 and D2 for the age-dynamic peak regions and their assigned genes. rho, Spearman’s correlation coefficient. (C) Genes associated with both age-dependent H3K4me3 and RNA expression changes tend to gain H3K4me3 marking in adult. Average plots show normalized H3K4me3 levels at L3, D2 and D12 for gene groups associated with only age-dependent H3K4me3 change (left), with both age-dependent H3K4me3 and RNA expression change (middle), and with only age-dependent RNA expression change (right). (D) The bar graph shows the percentages of age-dynamic H3K4me3 peaks that were accompanied by RNA expression change in the l to q clusters (with gene body H3K4me3) vs the other clusters (clusters described in Fig 2). Fisher’s exact test shows that the dynamic H3K4me3 peaks in clusters l to q are significantly more likely to be accompanied with age-dependent RNA expression change. (E) The lengths of the age-dynamic H3K4m3 peak are positively correlated with the fraction of the peak groups associated with RNA expression change. Age-dynamic H3K4me3 peaks assigned only to one gene are used for this analysis. The size of the dots indicates the number of peaks (also the number of genes) for each peak length range.

### Gene body H3K4me3 markings are enriched for age-dynamic H3K4me3 and gene expression changes

Although the age-dependent RNA expression changes and H3K4me3 changes were highly correlated, a substantial portion of the age-dynamic H3K4me3 peaks were associated with genes that showed no significant expression change with age based on our RNA-seq data (Fig 4A). Among the protein coding genes assigned to age-dynamic H3K4me3 peaks, 922 genes exhibited significant RNA expression change with age, whereas 1,650 showed stable expression (Fig 4A). To investigate the possible features that might distinguish these two groups, we first examined the average plots of H3K4me3 levels associated with the age-dynamic H3K4me3 peak groups that accompanied with or without RNA expression changes. The data showed that, compared to the genes associated with only H3K4me3 dynamics, the genes associated with age-dependent changes in both H3K4me3 and RNA expression were marked with relatively higher levels of gene body H3K4me3 coverage that were acquired during adult stage (Fig 4C).

To test the statistical significance of this observable difference, we took advantage of the 25 clusters we previously discussed and tested whether the gene body H3K4me3 clusters would be enriched for both dynamic H3K4me3 and gene expression changes with age. The results showed that, among the genes with dynamic H3K4me3, those in clusters 1-q, where H3K4me3 obviously marked gene body regions, were significantly enriched for genes with age-dependent RNA expression change compared with those in the other clusters (a-k, r-y)(Fig 4D). Furthermore, when individual clusters were evaluated, clusters k, 1, m and n, which are enriched for age-dynamic H3K4me3 peaks that cover gene body (Fig 2B), contained significantly higher percentage of genes that showed both H3K4me3 and RNA expression change (S3B Fig). Interestingly, cluster p, which contained genes marked by high levels H3K4me3 through gene body, while not enriched for age-dynamic H3K4me3 peaks (Fig 2B), nevertheless contained significantly higher fraction of genes that exhibited age-dependent change in both H3K4me3 and RNA expression (S3B Fig). We further detected a positive correlation between the length of the age-dynamic H3K4me3 peaks and the fraction of those peaks associated with genes expression change with age (Fig 4E). Taken together, the results suggested that broader regions of H3K4me3 marking, which tend to cover greater portion of the gene body and are mainly deposited in adult stage, are more prone to dynamic regulation with age and are also more likely to be accompanied by corresponding gene expression changes.

### Reduction of global H3K4me3 through *ash-2* RNAi results in altered expression of genes with adult-stage specific H3K4me3

We next wondered whether the specific features of H3K4me3 that we discovered to correlate with age-dependent regulation, including gene-body marking and adult-specific deposition, could have a role in regulating gene expression during adulthood. To test that, we analyzed previously published microarray data comparing the transcriptional profiles of D3 adult *glp-1* mutant worms with or without *ash-2* RNAi [35]. ASH-2 is a component of the H3K4me3 methyltransferase complex in *C. elegans* and *ash-2* RNAi initiated in larval stage 1 has been shown to cause a substantial global reduction of H3K4me3 in adult *glp-1* worms [35]. Interestingly, despite the global depletion of H3K4me3 levels, our analysis revealed only 831 and 981 protein-coding genes that showed increased or decreased mRNA expression respectively upon *ash-2* RNAi (S10 Table). Since H3K4me3 marking is well known to associate with actively expressed genes, the upregulated in mRNA expression following H3K4me3 depletion could reflect an indirect effect. In contrast, the downregulated in mRNA expression could reflect a more direct regulatory role of ASH-2 and H3K4me3 depletion.

We then intersected the candidate ASH-2-regulated genes with the gene lists we discussed earlier that associated with different features of H3K4me3 marking. Using GSEA, we found that the genes that exhibited downregulated mRNA expression when *ash-2* was knocked down were enriched for genes that exhibited greater H3K4me3 marking in D2 relative to L3, suggesting they were preferentially marked during adult stage (Fig 5A). The same relationship was demonstrated by average plots of the normalized H3K4me3 levels at the L3, D2, and D12 stages for the genes that were downregulated when *ash-2* was knocked down (Fig 5B). Interestingly, the genes that exhibited upregulated expression when *ash-2* was knocked down did not show this bias (Fig 5A and 5B). Moreover, the ASH-2-mediated upregulated genes appeared to be marked by higher levels of H3K4me3 starting at L3, whereas the ASH-2-mediated downregulated genes showed relatively low levels of H3K4me3 even during adulthood (Fig 5B). The results together indicated that when *ash-2* was depleted through larval and adult stages, which resulted in overall global reduction of H3K4me3 levels, a small specific set of genes responded with expression change. The mechanism for this selectivity is unclear, but the downregulated subgroup pointed to adult-stage specific H3K4me3 deposition to play an important role in maintaining the proper expression of genes during adulthood.

**Figure 5.**
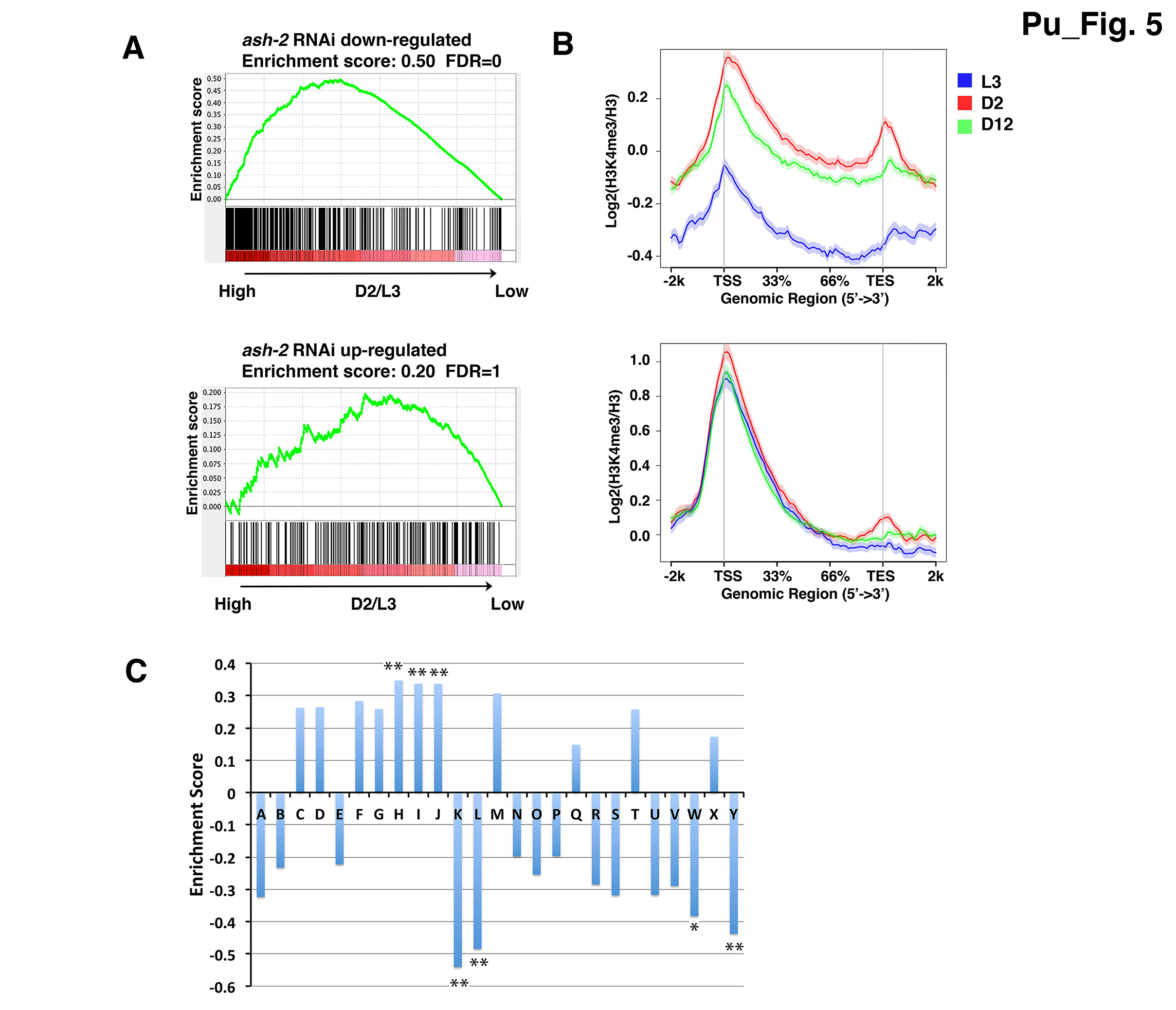
Genes that show down-regulated expression when *ash-2* was knocked down are more likely to be marked with H3K4me3 during adult stage. (A) For genes exhibiting significant down-regulated expression when *ash-2* was knocked down, GSEA revealed a significant enrichment for genes associated with higher D2 H3K4me3 marking compared to L3. For genes exhibiting significant up-regulated expression when *ash-2* was knocked down, GSEA revealed no bias. Genes with detectable expression from the published *ash-2* RNAi microarray experiment (ref) were ranked according to their H3K4me3 levels difference between D2 and L3 shown in Fig S2C. The genes that showed significant expression change comparing *ash-2* RNAi vs. control RNAi were first determined using limma in the R package (FDR <0.05), and the genes showing significantly decreased or increased expression were extracted for GSEA analysis. (B) Average plots show that the genes that exhibited significant down-regulated expression upon *ash-2* RNAi (top) were marked with higher levels of H3K4me3 at D2 compared to L3. For the genes that exhibited significant up-regulated expression upon *ash-2* RNAi (bottom), their average H3K4me3 levels did not change at the different developmental time points. (C) Clusters k, l, w and y were enriched for genes with down-regulated expression upon *ash-2* RNAi, and clusters h, i, j were enriched for genes with up-regulated expression upon *ash-2* RNAi. Genes in all of the 25 clusters were ranked according to their expression fold change as detected in the *ash-2* microarray experiment. GSEA was used to determine the enrichment for down-regulated or up-regulated genes in each cluster. (**), FDR<0.01, (*), FDR<0.05.

Interestingly, when we intersected the ASH-2 regulated genes with the 25 clusters previously defined (Fig 2), GSEA analysis showed that clusters k and l were significantly enriched for genes that became downregulated when *ash-2* was depleted. Since clusters k and 1 were also enriched for genes that showed age-dynamic H3K4me3 marking that tended to be adult deposited, the data together suggested that the age-dependent reduction of adult-deposited H3K4me3 likely contributes to the altered gene expression during aging.

### Genes associated with age-dynamic H3K4me3 and RNA expression change are enriched for stress response and fat metabolism functions

To gain possible biological insights into the genes associated with age-dynamic H3K4me3 and RNA expression change, we performed gene ontology (GO) analysis using DAVID (Sll Table). Among the protein coding genes associated with age-dependent H3K4me3 change, the functional clusters oxidation-reduction, fatty acid metabolism, pyridoxal phosphate and mitochondrion were most highly enriched for genes associated with decreased H3K4me3 with age. Similar functional clusters continued to be significantly overrepresented when only considering the genes associated with both H3K4me3 and RNA expression change with age. For the genes associated with increased H3K4me3 with age, the functional group “regulation of GTPase” was most highly enriched when RNA expression change was not considered. Interestingly, for genes associated with both increased H3K4me3 and RNA expression, stress response genes become highly overrepresented.

A link between stress response and aging has been well known, but the connection between fat metabolism and aging has been less well defined. A further inspection of the fat metabolism related genes revealed that ~47% of the annotated fat metabolism related genes show age-dependent dynamic changes in H3K4me3 marking (42 out of 89 genes). Among the 42 genes, 25 genes are located in clusters 1-n, with H3K4me3 mainly deposited on gene body regions. To investigate this further, we identified genes annotated to participate in fatty acid biosynthesis and fatty acid oxidation function based on WormBase, and compared them with the age-dependent H3K4me3 and RNA expression upregulated or downregulated gene set. This comparison revealed a strong overlap between fatty acid biosynthesis and fatty acid beta-oxidation related genes with H3K4me3 decreased and RNA downregulated genes. 35.1% of fatty acid biosynthesis related genes (13 genes out of 37 genes) and 23.5% of fatty acid oxidation related (8 genes out of 34 genes) are with reduced H3K4me3 and RNA abundance with age (Sll Table). The GO analysis suggested a possible deregulation of fat metabolism with aging. We monitored the fat content of *glp-1* mutants at various aging time points and confirmed that fat levels indeed decreased with age (S4C Fig).

## Discussion

In this study, we performed ChlP-seq analysis to identify dynamic changes of H3K4me3 with age in somatic cells of *C. elegans.* The data revealed that genome-wide H3K4me3 markings remain largely stable up until the D12 time point that we surveyed, but reproducible age-dependent changes in H3K4me3 marking are also readily detectable[45,46]. We found that ~30% of the H3K4me3 marked regions exhibit changes with age, and the dynamic changes generally happen in regions where H3K4me3 markings are relatively lower. In contrast, the regions marked by high levels of H3K4me3 remain largely stable with age. It is important to note that our experimental design, which used whole worm extracts, cannot distinguish regions marked by low levels of H3K4me3 in many cells vs. regions marked by high levels of H3 K4me3 in a small subset of cells.

Our analyses revealed two major patterns of H3K4me3 marking: The canonical pattern where H3K4me3 marks concentrate around transcriptional start sites with a bias towards the 5’ promoter regions [23], and the atypical pattern where H3K4me3 marks span gene body regions. For the gene body spanning H3K4me3, we also observed either high or low levels of H3K4me3 markings. Interestingly, our data indicated that the canonical pattern of H3K4me3, as well as the gene body H3K4me3 with high levels of marking, are largely established by the L3 stage, and these H3K4me3 markings generally remain stable through the aging time points that we surveyed. In contrast, the weak or moderate levels of gene body H3K4me3 markings are often acquired during adulthood and they exhibit dynamic changes with age. These adult-stage specific H3K4me3 marked regions could reflect unique activities of histone modification enzymes in post-mitotic cells, but also could reflect marking acquired in specific subset of cells during adulthood.

Histone modification marks are well known to act in concert to maintain local chromatin environment, we next compared the H3K4me3 profiles reported here with the H3K36me3 profiles we previously published [44]. As has been previously described, we identified regions co-occupied by both H3K4me3 and H3K36me3 around TSS and spanning gene body [23]. This pattern of co-occupancy is evident starting in worms at the L3 stage (S2A Fig). Interestingly, for the adult-specific H3K4me3 regions, which exhibit greater dynamics with age, we observed low or undetectable levels of H3K36me3 (S2A Fig). These genes are generally actively expressed in adults but are more likely to show expression changes with age (S2A Fig). This observation is exactly consistent with our previous finding that H3K36me3 marking is necessary for maintaining gene expression stability during aging and genes marked by low or undetectable levels of H3K36me3 show greater age-dependent RNA expression dynamics [44,47].

Since the age-dynamic H3K4me3 regions appear to be adult-stage specific, we also examined the genome-wide pattern of the histone variant H3.3, HIS-72 in *C. elegans* somatic cells. Histone variant H3.3 is the major source for H3 turnover in post-mitotic cells and is thought to contribute to enhanced epigenetic plasticity and nucleosome dynamics [48]. We observed that the age-dynamic H3K4me3 regions usually have higher levels of H3.3 compared to the stable H3K4me3 regions. In summary, we found that most of the H3K4me3 peaks that change with age are deposited in adult stages, and embedded in a chromatin environment with low levels of H3K36me3 and high levels of histone variant H3.3. These results pointed to interesting temporal and spatial regulations of H3K4me3 depositions with important consequence on H3K4me3 stability with age. Future investigation of the mechanisms of this regulation will provide important new insights into the relationship between chromatin marks and gene expression regulation.

Our RNA-seq analysis revealed that ~7% of the protein-coding genes exhibit altered expression with age. Even though H3K4me3 is thought to mark most of the actively expressed genes, we found that ~30% of the genes that showed significant expression change between D2 and D12 were not detectably marked by H3K4me3 at either time points (S5 Table), which could be due to low or cell type specific expression. It is interesting to note that ~70% of the genes that showed H3K4me3 changes are not accompanied by RNA changes. And ~50% of the genes that showed RNA changes are not accompanied by H3K4me3 changes. This observation is consistent with the current thinking that H3K4me3 marking is not instructive for gene expression, but rather represents a mark of transcriptional history. Following this thinking, changes in H3K4me3 would not be sufficient to induce RNA expression changes, and RNA expression changes do not necessarily have to be accompanied by H3K4me3 changes, depending on how quickly transcriptional “memory” follows transcriptional changes. Moreover, some of the changes detected in our RNA-seq data may not reflect transcriptional changes, but rather could be due to altered RNA stability.

An important finding from our study is that there is a subset of genes that show consistent and significant changes in both H3K4me3 and RNA expression levels with age. The H3K4me3 modification in this gene group showed a bias of adult stage deposition on gene body regions (Fig 4C and S3A Fig). Interestingly, in examining the RNA profiling data associated with a global reduction of H3K4me3 with *ash-2* RNAi, we revealed that only a small subset of genes responded with decreased RNA expression. Since H3K4me3 is a histone mark that associates with active gene expression, if it had a regulatory role in promoting gene expression, its loss would be expected to result in decreased gene expression. Intriguingly, the subset of genes that decrease expression when H3K4me3 levels are reduced have a high tendency to be genes that normally acquire gene-body H3K4me3 marking during adulthood. While this finding cannot be used as definitive evidence of cause- and-effect relationship between the H3K4me3 and RNA expression change with age, the data are certainly consistent with the model that altered H3K4me3 levels can have a regulatory role on the RNA expression of the specific subset of genes whose gene-body H3K4me3 mark are deposited during adult stage. Studies in multiple organisms have shown that gene body H3K4me3 is associated with RNA polymerase elongation [34,49,50]. It is possible that RNA polymerase elongation efficiency alters with aging, which could contribute to the age-dependent RNA expression changes observed for the subset of genes.

Our GO term analysis suggested that the genes that showed age-dynamic H3K4me3 and RNA expression change are enriched for a number of different functional groups, including fat metabolism. Interestingly, recent findings suggested that the elevated expression of the fatty acid desaturase enzymes FAT-5 and FAT-7 is essential for the lifespan extension caused by global inactivation of the H3K4me3 methyltransferase complex in wild-type reproductive worms [37]. However, when we inspected our data from the germlineless *glp-1* mutant worms, we noted that *fat-5* is marked with high levels of H3K4me3 on gene body (cluster o) and its H3K4me3 marking and RNA expression do not show age-dependent change. In addition, even though *fat-7* is actively expressed, it is not associated with detectable H3K4me3 marking. Therefore, our data indicated that *fat-5* and *fat-7* are not among the small subset of genes whose H3K4me3 and RNA expression show age-dynamic regulation. This result is not surprising since global reduction of H3K4me3 via inactivation of the methyltransferase is thought to act through the germline to regulate specific aspects of fat metabolism and to modulate lifespan. We speculate that in the somatic cells of *C. elegans,* a small subset of genes of particular biological importance, particularly in fat metabolism and stress resistance, acquire adult-specific H3K4me3 markings that span gene body and are dynamically regulated through aging. This age-dynamic H3K4me3 profile in turn regulate RNA expression change and likely contribute to specific physiological changes that accompany aging (Fig 6).

**Figure 6.**
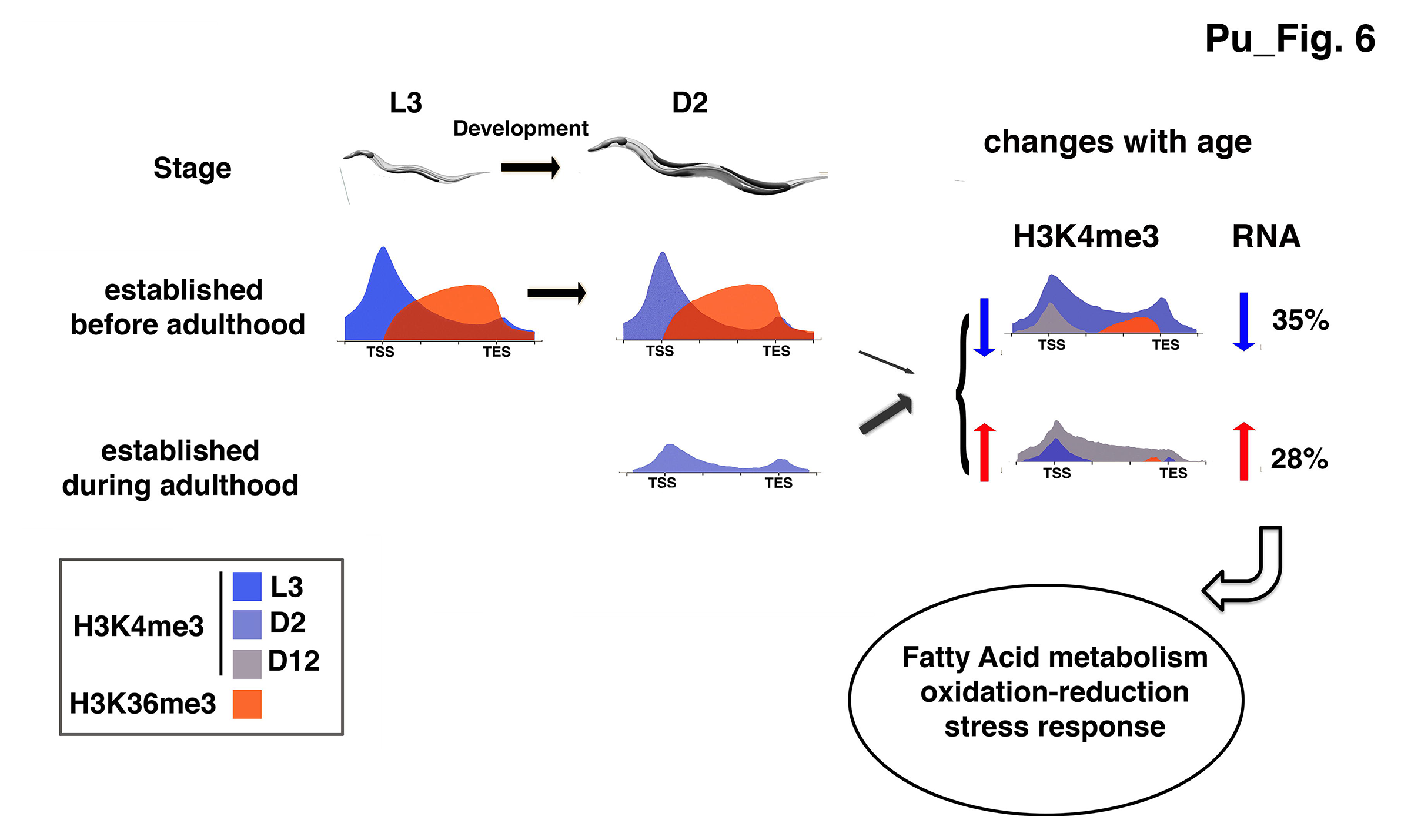
H3K4me3 marking established during adulthood are more likely to be dynamically regulated with age and be accompanied by corresponding RNA expression. Our data revealed that the H3K4me3 markings established before adulthood generally remain stable with age. These H3K4me3 marks have the canonical pattern of accumulating to high levels around TSSs, with H3K36me3 marking the corresponding gene bodies. In contrast, the H3K4me3 markings established in adult stages tend to be of lower levels and distribute more evenly into gene bodies that are marked with very low or non-detectable levels of H3K36me3. The width of purple arrow indicates the proportion of genes with significant age-dependent H3K4me3 change for each H3K4me3 marking pattern. For the genes associated with significant increased (blue) or decreased (red) H3K4me3 levels with age, ~35% or 28% are accompanied with corresponding RNA expression change.

## Methods

### *C. elegans* strain growth and harvesting

*C. elegans* strain *glp-1(e2141)* is cultured as previously[44]. *glp-1(e2141)* stocks were kept at 16°C and grown under standard growth conditions[51]. For ChlP-seq and RNA-seq experiments, embryos prepared from 16°C *glp-1(e2141)* stocks by bleaching were hatched and cultured at 25°C with 3000 embryos per 15 cm of nematode growth medium (NGM) plate seeded with 1.5 mL of concentrated E. coli OP50 (30× overnight culture) with 50 μg/mL carbenicillin and 15 μg/mL tetracycline. For D12 samples, worms were re fed once on D4. Adult worms at the D2 and D12 stages were washed with ice-cold M9 three times. Worm pellets were stored at −80°C before ChIP and RNA extraction.

### ChIP and ChlP-seq library preparation

ChIP was performed as described [52,53]. Worm pellet was ground with a mortar and pestle and cross-linked with 1% formaldehyde in PBS for 10 min at room temperature. Worm fragments were collected by spinning at 3000g for 5 min and resuspended in FA buffer followed by sonication with Bioruptor to generate chromatin fragments with major DNA length around 200 bp. Chromatin extract was incubated with H3 antibody (rabbit; Abcam, abl791), H3K4me3 antibody (rabbit; Abeam ab9050) overnight at 4°C. Antibodies used were prescreened for specificity using dot blots. The optimal amounts of antibodies used were determined by titration in a preliminary experiment with ChlP-qPCR. Precipitated DNA (10-15 ng) from each sample was used for Illumina sequencing library preparation. DNA from ChIP was first end-repaired to generate a blunt end followed by adding single adenine base for adaptor ligation. The ligation product with adaptor was size-selected and amplified by PCR with primers targeting the adaptor. Up to 12 samples were multiplexed in one lane for single-end 50-nt Illumina HiSeq sequencing or single-end 75-nt Illumina NextSeq500. Raw ChlP-seq data have been deposited at Gene Expression Omnibus (GSE101964).

### ChlP-seq data analysis

Sequencing reads were filtered by FASTX_toolkit (-q 20 -p 80) and mapped to *C. elegans* reference genome (celO) by bowtie2 with the default setting [54]. Mapped reads were extracted and PCR duplicates were removed with samtools [55].

To compare the independent replicates of ChIP experiment, the mapped Illumina reads were counted in 2 kb sliding windows across the *C. elegan* genome in each experiment separately. The tag counts were normalized by the total number of aligned reads of each experiment, including the control experiments of H3, then multiply that by a million to get CPM. Lastly, log2 transformed CPM were calculated from treatment experiments and control experiments separately, and the control values were subtracted by corresponding treatments ones, which used for calculating Pearson Correlation. Only the replicates, with values higher than 0.8, used for further analyses. All statistical analysis was performed in R environment.

For H3K4me3 peak calling, mapped reads from biological replicates were merged for peak identification with MACS2 (2.1.0)[56]. Narrow and broad peaks were called by MACS2 with default settings for narrow peak calling or broad peak calling. Peaks were further verified with 3 independent replicates by using GLM method in R package. Verified peaks were used for time point differential analysis by Diffbind (version 3.2.1) to identify dynamic peaks significantly changed between time points. The identified dynamic peaks that were identified from narrow peak calling or broad peak calling were merged by lbp over-lapping and were further filtered with normalized histone modification levels and fold changes. H3K4me3 and H3 signals in each peak were calculated with Homer normalized to total unique mapped reads in each library. H3K4me3 levels were further normalized with H3. Peaks with ratio of H3K4me3 and H3 higher than 1.0 in all three replicates at either time point were kept. Further, the peaks with normalized H3K4me3 levels increased or decreased by at least 30% during aging were kept for downstream analysis.

H3K4me3 peaks were assigned to the closest genes annotated in WBCel235 by Bedtools. The assignments with H3K4me3 peaks located downstream of genes without overlapping were removed.

H3K4me3 profiles of protein coding genes associated with H3K4me3 peaks at D2 and/or D12 adult were clustered into 25 clusters by ngsplot [57] with -GO km setting with -KNC 25. The difference of H3K4me3 peaks between D2 and L3 were ranked by using ngsplot with -GO diff setting.

### GSEA analysis

For GSEA analysis, GseaPreranked tool was used with the pre-ranked gene list. The ranks of peaks displayed in heatmaps were assigned to the associated genes. Genes associated both with strong D2 and strong L3 stage H3K4me3 signals were exclude from the analysis. If more than one rank were assigned to one gene, the higher rank was kept for GSEA analysis.

### RNA-seq library preparation and data analysis

Total RNA was extracted from worms harvested at the same stages as ChlP-seq sample preparation for *glp-1(e2141)* worms using TRI reagent (Molecular Research Center). Total RNA were used for library preparation with nugen kit (). 4 samples were multiplexed in one lane for single-end 100nt Illumina HiSeq sequencing. Raw RNA-seq data have been deposited at GEO (GSE101964).

For data analysis, tRNA and rRNA reads were first filtered out using Bowtie2, and the remaining reads were further aligned to WBcel235 transcript annotation by TopHat2 with no novel junctions allowed (Trapnell et al. 2009, 2012). The accepted aligned reads with a maximum of two mismatches were kept for differential expression analysis using edgeR. 46,590 genes were actively expressed with gene expression levels between 0.06 CPM and 11333.82 CPM. FDR cutoff of 0.05 was set for identifying differentially expressed genes.

### Oil-Red-O (ORO) staining of worms

Oil-Red-O staining was performed as described [58]. Worms were washed with M9 and stored at −80°C. For staining, worms were first fixed with 60% isopropanol and stained with freshly prepared Oil Red 0 working solution at 25 °C in a wet chamber for 6-18 hours. The Oil Red 0 working solution was removed and worms were then washed with 0.01% Triton X-100 in S buffer and kept in this solution at 4 °C before imagining.

### Availability of data

ChlP-seq and ribo-RNAseq data from this study have been submitted to the NCBI Gene Expression Omnibus (GEO; http://www.ncbi.nlm.nih.g0v/ge0/) under accession number GSE101964.

## Acknowledgments

We thank Dr. Charles Danko (Cornell University) for insightful discussion and manuscript reading. We thank Dr. Jen Grenier (Cornell University) for help with ribo-RNAseq sequencing data analysis, Dr. Veerle Rottiers (Cornell University) for help with oil-o-red staining, Dr. Françoise Vermeylen (Cornell Statistical Consulting Unit) for help with statistical analysis. Some strains were provided by the CGC, which is funded by NIH Office of Research Infrastructure Programs (P40 OD010440).

## Supporting Information Legends

### Supplemental Figures

**SI Figure. Related to Figure 1.**
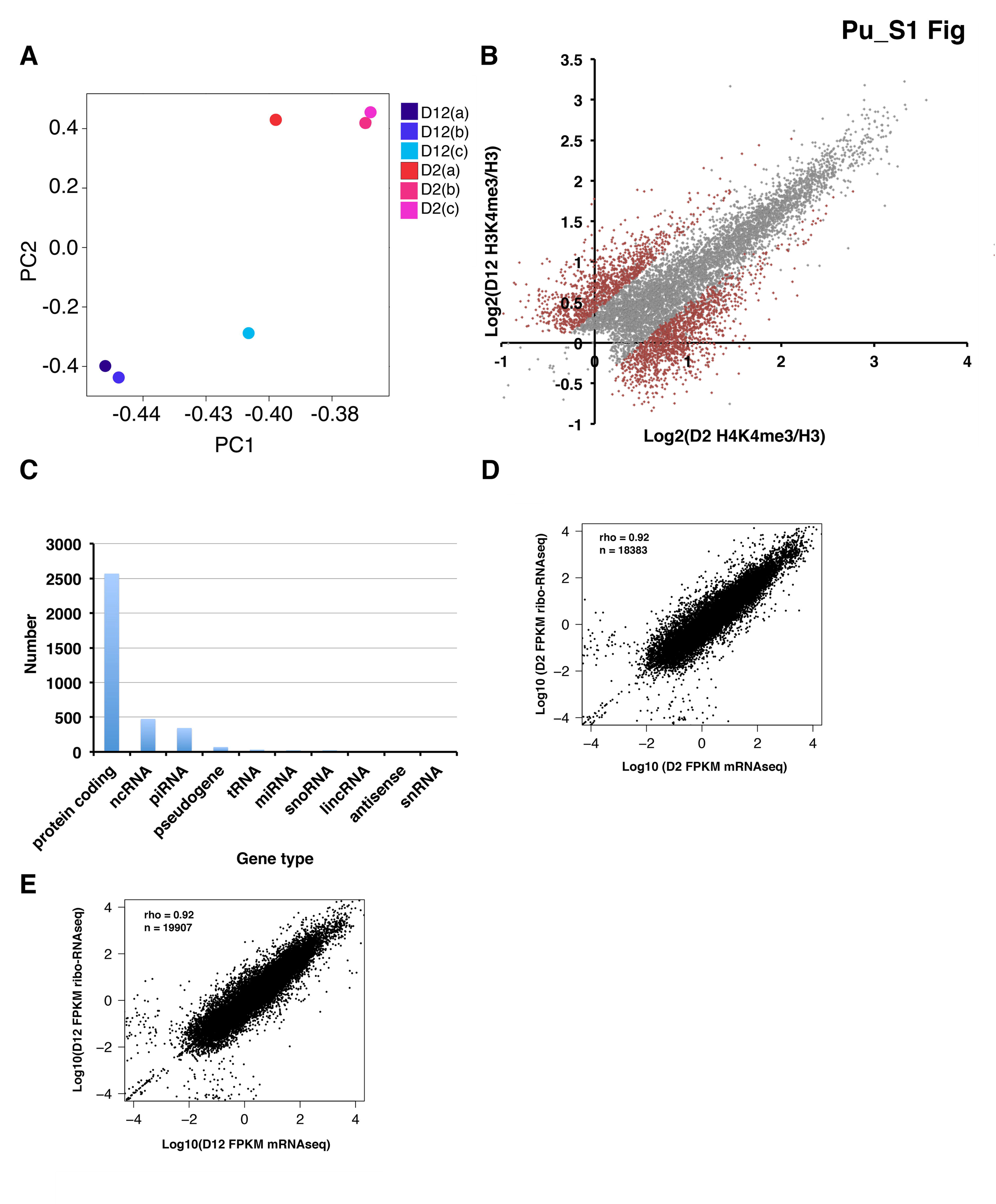
(A) PCA plot showing normalized H3K4me3 data from three biological replicates. The H3K4me3 peaks in D2 and D12 were identified by the MACS2 broad peak calling method. (B) Scatter plot showing normalized H3K4me3 levels at D2 and D12 for stable (grey) or dynamic (brown) H3K4me3 peaks. The plot shows the average normalized H3K4me3 signals from three biological replicates calculated using Homer. (C) Each dynamic H3K4me3 peak was assigned to its closest gene as annotated in WBcel235. Assigned gene numbers of different gene types are shown. (D) and (E) Comparison of mRNA-seq data and ribo-RNAseq data from D2 (D) or D12 (E) *glp-1 (e2141)* worms. The loglO (FPKM) values of mapped genes in both experiments were compared.

**S2 Figure. Related to Figure 3.**
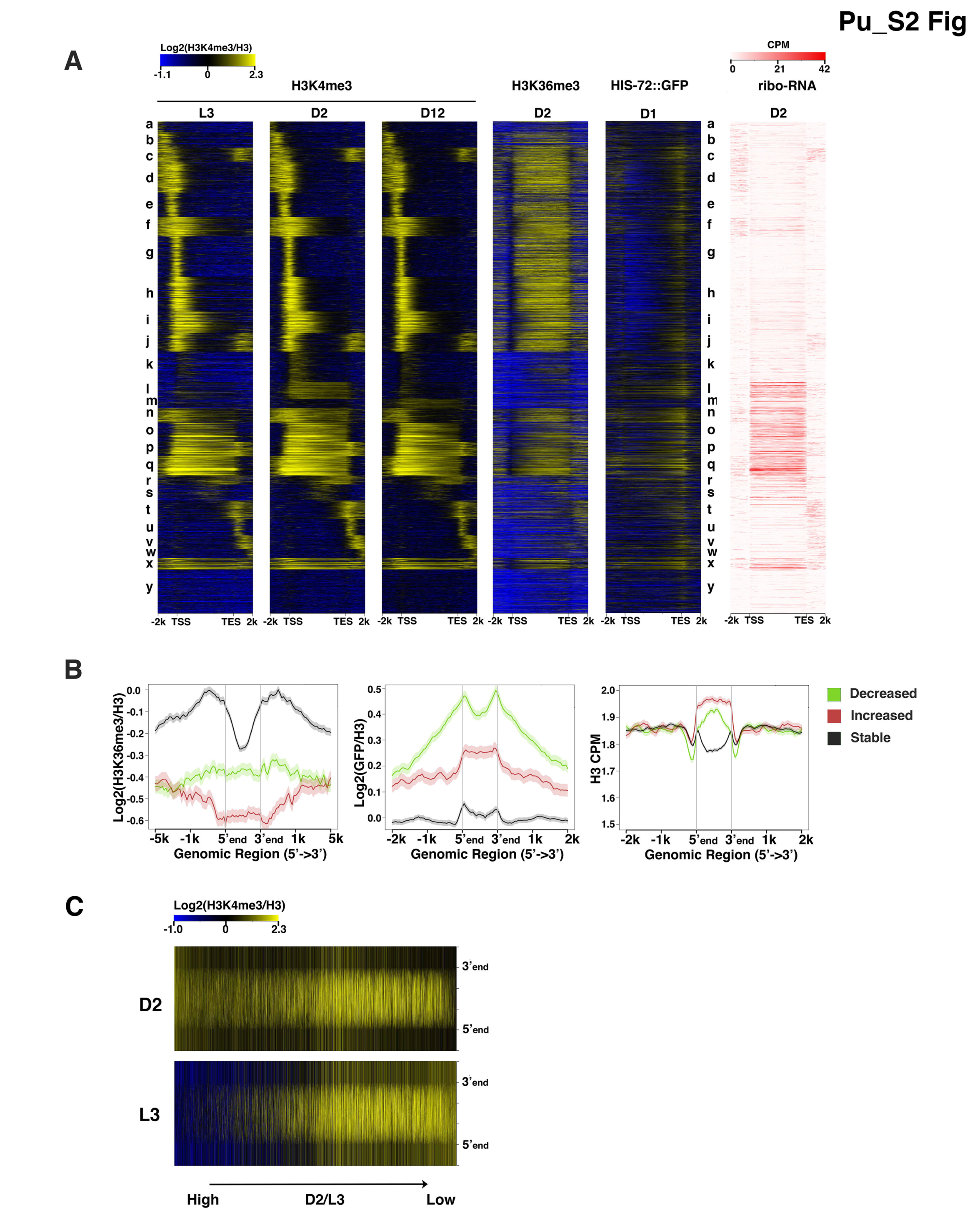
(A) Heatmap showing the normalized ChlP-seq signals of H3K4me3, H3K36me3, HIS-72::GFP and RNA abundance at the indicated stages in the 25 clusters generated by k-mean clustering in Fig 2A. (B) Average plots show normalized H3K36me3, HIS-72::GFP and H3 signals within and surrounding H3K4me3 peaks that decreased (green), increased (red) or remained stable with age (black). (C) Heatmaps show the H3K4me3 levels of D2 stage H3K4me3 peaks at D2 (top) and L3 (bottom) stages. H3K4me3 peaks were ranked according to the ratio of H3K4me3 levels at D2 to that of L3. Peaks with higher D2 H3K4me3 levels were put on the left side of the heatmap.

**S3 Figure. Related to Figure 4.**
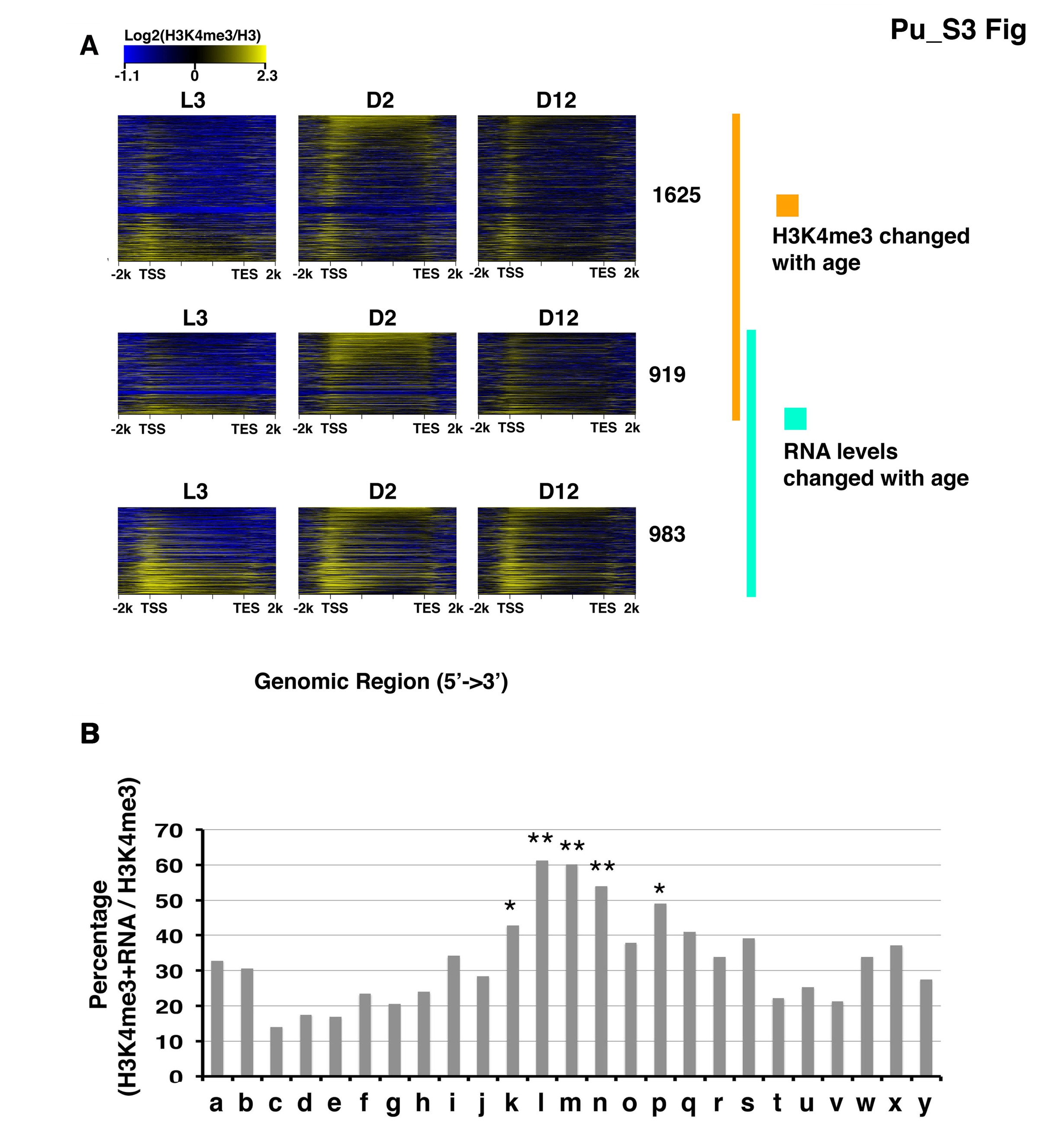
(A) Heatmaps showing normalized H3K4me3 signals at L3, D2 and D12 of the indicated gene groups with or without age-dependent H3K4me3 and/or RNA expression changes. The genes in each heatmap panel were ranked according to the ratio of H3K4me3 levels at L2 to that of L3. Peaks with higher ratios were placed at the top of the heatmaps. (B) The bar chart shows the percentage of genes associated with age-dynamic H3K4me3 that also exhibited age-dependent RNA expression change in each of the 25 clusters shown in Fig 2A. Clusters k, l, m, n and p are with significantly higher occurrence of age-dependent H3K4me3 changes that were accompanied by RNA expression changes.

**S4 Figure. Related to Sll Table.**
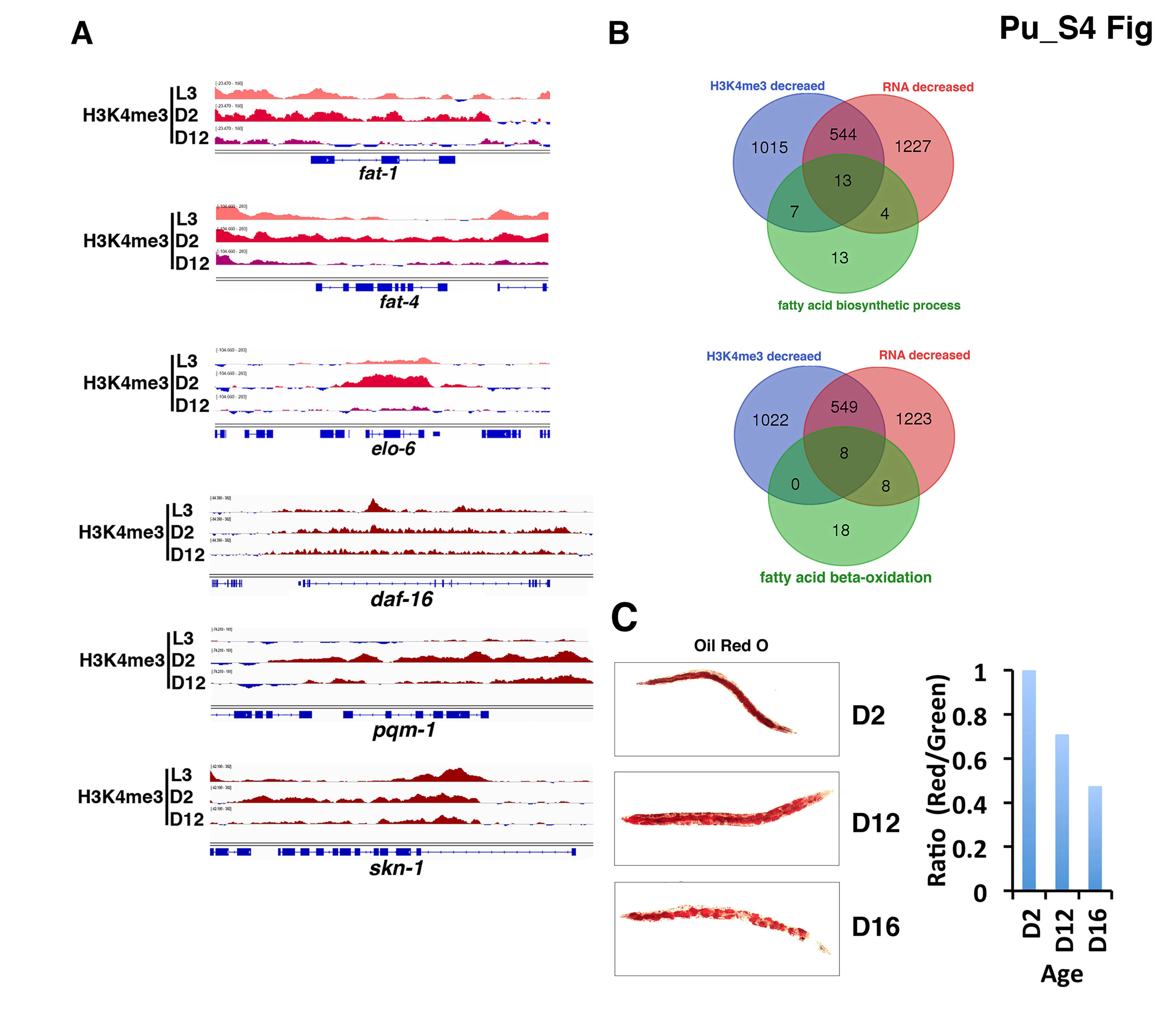
(A) Representative genome browser views of H3K4me3 marking around fat metabolism genes and lifespan regulators at the indicated stages. (B) A significant fraction of the genes annotated in the fatty acid biosynthetic (top) or fatty acid beta-oxidation (bottom) pathways showed decreased RNA expression and/or H3K4me3 levels with age. Red: Genes with decreased RNA expression with age. Blue: Genes with decreased H3K4me3 levels with age. Green: Genes annotated to participate in fatty acid biosynthetic or beta-oxidation processes. Gene numbers of each group are shown in the venn diagrams. (C) Oil-Red-O (0R0) staining of worms at D2, D12 and D16 aging time points (right). Quantification of the 0R0 signals (left) was done by normalizing the total signal counts in the red channel with those in the green channel.

### Supplemental Tables

**S1 Table: Summary of ChlP-seq and RNA-seq**

**S2 Table: MACS2 identified H3K4me3 peaks**

**S3 Table: Diffbind identified age-dynamic H3K4me3 peaks**

**S4 Table: Dynamic H3K4me3 peaks associated genes**

**S5 Table: 25 clusters of protein coding genes associated with H3K4me3 peaks.**

**S6 Table: H3K4me3 peaks in *glp-l* at L3 stage.**

**S7 Table: Diffbind identified differential H3K4me3 peaks between L3 and D2 adult.**

**S8 Table: edgeR result of ribo-RNAseq**

**S9 Table: Correlation between H3K4me3 changes and RNA expression changes of genes associated with dynamic H3K4me3 peaks**

**S10 Table: Genes significantly changed expression upon *ash-2* RNAi at D3 adult *glp-l.***

**S11 Table: Functional annotation of genes with H3K4me3 change and RNA level change.**

